# Orsay virus infection is associated with reduced locomotion and alters cue-dependent food choice in *Caenorhabditis elegans*

**DOI:** 10.64898/2026.11.04.764174

**Authors:** Dominik Herek, Esmeralda G. Legarda, Simone Carrara, Ana García-Ruiz, Beren İstek, Izan Melero, Paula Pons, Ana Villena-Giménez, Victoria G. Castiglioni, Santiago F. Elena

## Abstract

Behavioral responses are an important component of host defense; however, most evidence in *Caenorhabditis elegans* comes from bacterial pathogens. Here, we tested whether infection with Orsay virus (OrV) alters host dispersal and food choice and whether these responses involve the RNA-processing factor EOL-1. When measuring distance travelled over a certain period of time in control and infected wild-type animals, we observed that OrV infection was associated with lower locomotion. Moreover, when presenting infected and control animals with food choices, we observed that infected animals preferred OrV infected food sources only when the alternative was a pure food source, preferring a non-inoculated food source when both control and viral food sources contained a *C. elegans* filtrate. Moreover, loss of EOL-1 altered baseline locomotion and attenuated the infection-associated reduction detected in the earlier motility assay, while also modifying the food choice response, suggesting that *eol-1* contributes to the motility and behavioral changes induced by OrV infection. Collectively, these results support an overall association between OrV infection and reduced host dispersal, whereas its effect on food choice depends on the sensory context. These findings extend infection-associated behavioral responses to a natural viral pathogen and identify *eol-1* as a modulator of their expression.

**IMPORTANCE:** This study advances our understanding of the interplay between viruses and host behavior by demonstrating that natural infection with OrV is associated with reduced motility and alters food choice in *C. elegans* in a cue context-dependent manner. In addition, it identifies *eol-1* as a genetic modulator of these responses, linking infection status to sensory processing and behavior. Taken together, these findings extend the concept of behavioral immunity beyond bacterial pathogens and establish the *C. elegans*-OrV system as a valuable model for investigating the neurogenetic basis of behavioral changes induced by viral infections.

## Main

*Caenorhabditis elegans* continuously interacts with complex microbial communities that include both beneficial and pathogenic organisms. In this ecological context, behavior constitutes a critical first line of defense against infection, complementing physiological and immune responses (1–3). Given that the immune responses are conserved, and its nervous system is well characterized, *C. elegans* has emerged as a powerful model system to study the interplay between the host, microbes, and behavior (4–8). As a result, a growing body of work has demonstrated that microbial exposure can profoundly influence *C. elegans* behavioral states, thereby shaping locomotion, feeding, and sensory decisions (9–12).

One of the best-established behavioral responses to infection in *C. elegans* is the emergence of so-called “sickness behaviors”, which include reduced locomotion, decreased feeding, and increased quiescence (13,14). For instance, chronic infection with the bacterial pathogen *Pseudomonas aeruginosa* induces marked reductions in exploratory behavior and promotes quiescent states mediated by neuro-modulatory circuits linked to stress and satiety (14). Similarly, infection with *Serratia marcescens* leads to locomotor quiescence whose magnitude scales with infection severity (15–17). In these systems, reduced mobility is a conserved and regulated response to infection, rather than a simple byproduct of pathogenic damage. Mechanistically, such behavioral changes arise from the tight coupling between peripheral infection signals and neural circuits, involving specific neurons and neuropeptides that modulate global behavioral states (5,14,18).

In addition to general sickness responses, *C. elegans* exhibits sophisticated behavioral adaptations that allow it to detect and selectively avoid harmful microbes. Through chemosensory perception and learning, animals can distinguish between nutritive and pathogenic bacteria and modify their behavior accordingly, often transitioning from initial attraction to learned avoidance following exposure to pathogenic cues (6,10,19). These avoidance behaviors rely on the integration of external sensory information, such as small RNAs produced by pathogens (19,20), and internal physiological states and are mediated by defined neural circuits that coordinate motor output with infection status (21–23). Such behavioral plasticity is considered a key component of the “behavioral immune system”, enabling animals to minimize exposure to pathogens and mitigate infection risk (5,14,21).

Importantly, behavioral responses to infection are tightly linked to neuroimmune signaling pathways. Neurotransmitters and neuromodulators such as serotonin, dopamine and neuropeptides regulate both immune defenses and behavioral outputs, highlighting the bidirectional crosstalk between the nervous and immune systems (9,11,18,23–28). This integration ensures that behavioral responses are appropriately tuned to infection status and physiological needs.

Although most of our understanding of infection-induced behavior in *C. elegans* derives from studies with bacterial pathogens, little is known about how viral infections influence host behavior. The Orsay virus (OrV), the only known natural viral pathogen of *C. elegans*, has been widely used to characterize antiviral immunity and host transcriptional responses (13,29). Importantly, naturally associated bacteria can also profoundly modulate susceptibility to OrV infection, highlighting the ecological relevance of interactions between the host, its microbial environment, and viral pathogens (30). However, emerging evidence indicates that OrV infection can also influence host behavior. Given that responses to OrV share common features with other biotic stressors, it is plausible that viral infection may also induce behavioral changes analogous to those observed with bacterial pathogens. In particular, viral infection has been shown to modulate male sexual behavior (31), while other studies have demonstrated that OrV infection can induce quiescence, suggesting that behavioral changes may constitute a key component of the host response to viral infection (13).

In this study, we investigated the effect of OrV infection on locomotion and food choice in *C. elegans*. Using four complementary population- and individual-level assays, we quantified infection-associated changes in locomotion and dispersal. We also tested whether infection altered food choice behavior, potentially through changes in sensory decision-making. To separate locomotor effects from food preference, we analyzed two components of the food-choice assay separately: (*i*) the probability of reaching either food patch during the observation period and, among animals recovered from a patch, (*ii*) the probability of selecting the OrV-containing preparation. We further examined the mechanistic basis of these behavioral changes by assessing the contribution of *enhanced olfactory learning-1* (*eol-1*), which is upregulated during OrV infection and has been implicated in sensory processing (32). EOL-1 is homologous to vertebrate DXO/Dom3Z, a conserved decapping exoribonuclease involved in pre-mRNA quality control (33). Accordingly, it is predicted to possess 5’→3’ exonuclease and mRNA 5’-diphosphatase activities (34). In *C. elegans*, EOL-1 localizes to the nucleus and cytoplasm and is expressed in several cell types, including the URX, AVF, and PQR neurons, other oxygen-sensing neurons, and reproductive tissues. Its activity in URX neurons inhibits olfactory learning, including learned aversion to *P. aeruginosa* (33), and its functional conservation is demonstrated by the ability of mouse Dom3Z to substitute for EOL-1 in *C. elegans* (33,35). EOL-1 is also necessary to induce the RNA interference pathway upon mitochondrial dysfunction (36). By comparing the surrogate wild-type (WT) strain ERT54 with *eol-1* mutants, our design allowed us to identify a reproducible effect of infection on dispersal, distinguish it from context-dependent changes in food selection, and evaluate the genetic contribution of *eol-1* without interpreting failure to reach a food patch as sensory indecision.

## RESULTS

### OrV infection was associated with reduced *C. elegans* locomotion across assays

Motility experiments were performed in two different manners (Fig. 1A). In both cases, animals inoculated after hatching were placed in the center of the plates and observed after different amounts of time, therefore raising different stages. In the first experiment (AGR.motility, Fig. 1B), the position of the animals was scored in a set of concentric circles, whereas in the other three experiments (PP.motility, Fig. 1C; SC.motility, Fig. 1D; Bİ.motility, Fig. 1E), the plates were photographed, and the path traveled by the animals was measured. In our first motility assay (Fig. 1B, AGR.motility), infection status on young adult (YA) animals had a significant effect on the dispersal distance (χ^2^ = 13.677, 1 d.f., *P* < 0.001). These infected animals were significantly more likely to occur in lower distance categories than noninfected animals, indicating reduced dispersal during the 1 h observation period. The estimated effect size from the CLMM was −0.762 ±0.206 (±1 SE), corresponding to an odds ratio (OR) of 0.47. Thus, infected animals had approximately 53% lower odds of reaching higher distance categories than noninfected animals. Although the distance from the plate center remained the primary determinant of animal distribution, infection significantly altered this spatial pattern under the conditions tested.

**FIG 1.**
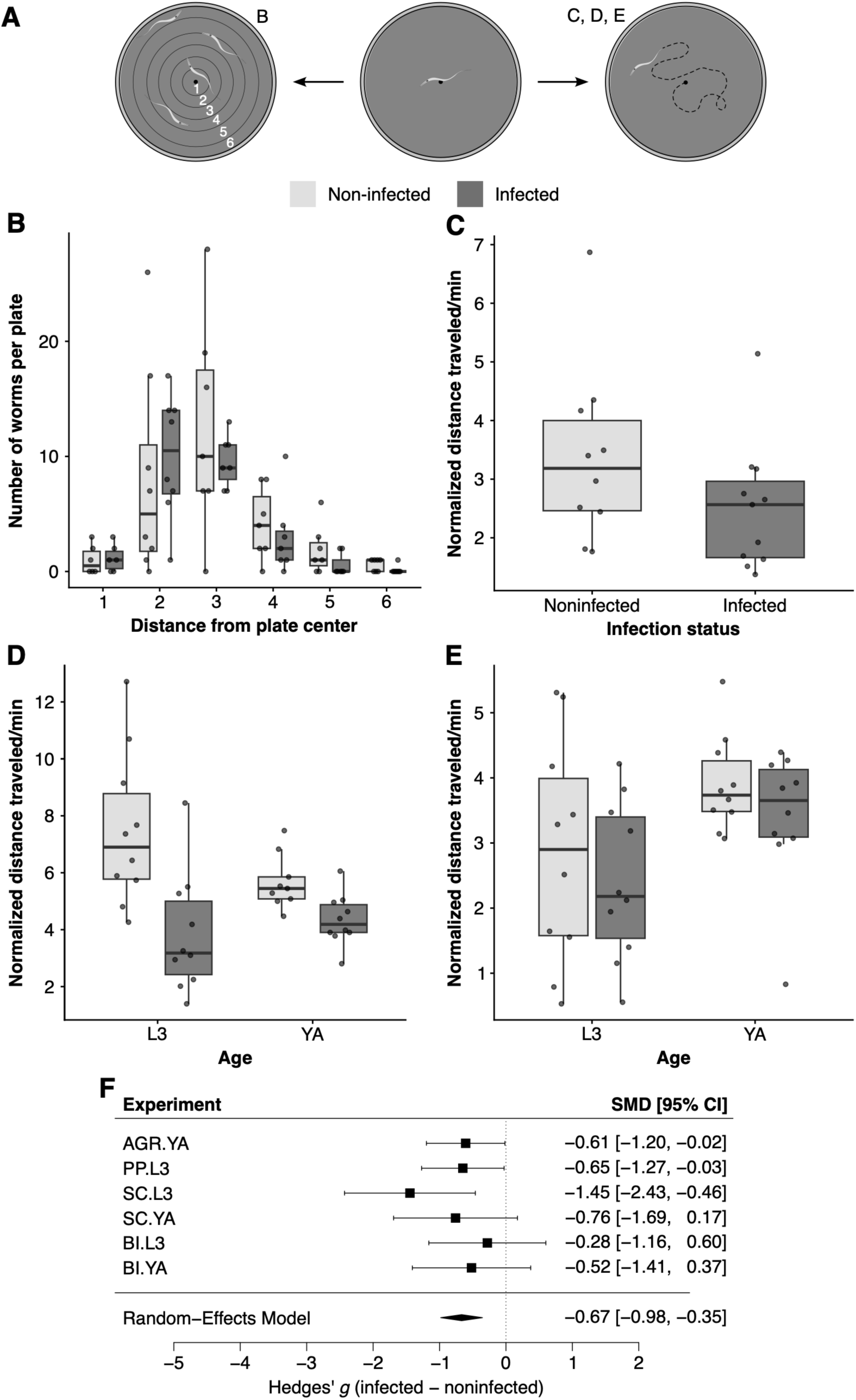
Effect of OrV infection on *C. elegans* motility. **(A)** Schematic of the different experimental designs used to measure motility. The corresponding figure panels are listed next to each plate diagram. **(B)** Experiment AGR.motility; spatial distribution of noninfected and infected YA animals relative to the plate center after 1 h. Each point represents an independent plate replicate. **(C)** Experiment PP.motility; body-length normalized distance traveled per minute (BLN/min) by noninfected and infected L3 individuals after 105 min. **(D)** Experiment SC.motility; BLN/min by noninfected and infected individuals from developmental stages L3 and YA after 30 min. **(E)** Experiment Bİ.motility; BLN/min by noninfected and infected L3 and YA individuals after 65 min. In panels (B) - (E), boxes depict the IQR, horizontal lines indicate medians, whiskers extend to 1.5×IQR, and points represent individual measurements. **(F)** Multilevel meta-analysis of the effect of OrV infection animal motility across six experiments. Forest plot showing standardized mean differences (Hedges’ *g*) for infected *vs*. noninfected animals in each experiment. Squares represent the effect size estimate for each experiment, with horizontal lines indicating 95% CI. The diamond represents the pooled effect estimated using REML, with its width corresponding to the 95% CI.

Fig. 1C shows the results from the second motility assay (PP.motility), which was performed on third-stage larvae (L3). In this assay, OrV infection had a significant negative effect on locomotion (χ^2^ = 3.495, 1 d.f., 1-tailed *P* = 0.031). On average, the movement rate of infected animals was 25.7% lower than that of noninfected animals, reaching 2.5 ±0.3 body-length equivalents (BLE)/min compared with 3.4 ±0.5 BLE/min in noninfected animals.

Fig. 1D shows the results of the third experiment (SC.motility). This experiment included observations of two developmental stages, L3 larvae and YA. Overall, infection status had a highly significant effect on motility (*F*_1,34_ = 19.627, *P* < 0.001). In contrast, developmental stage had no significant main effect (*F*_1,34_ = 1.353, *P* = 0.253), indicating that L3 and YA animals traveled similar BLE distances over the 30 min observation period. However, the interaction between infection status and developmental stage was significant (*F*_1,34_ = 4.294, *P* = 0.046), indicating that the effect of infection depended on developmental stage. Infection reduced locomotion more strongly in L3 larvae, in which the movement rate declined by 48.7% (3.8 ±0.7 BLE/min in infected animals *vs* 7.5 ±0.8 BLE/min in noninfected), than in YA animals, where locomotion was reduced by 23.2% (4.3 ±0.3 BLE/min in infected animals *vs* 5.7 ±0.3 BLE/min in noninfected animals).

The fourth experiment (Bİ.motility) followed the same design as the previous assay, but locomotion was measured in a 65 min observation period (Fig. 1E). Developmental stage was the only factor with a significant overall effect on motility (*F*_1,35_ = 7.664, *P* = 0.009). In contrast, OrV exposure had a negative, but not significant, effect on motility in both L3 and YA animals. Among L3, infection reduced the movement rate by 15.4% (2.4 ±0.4 BLE/min in infected animals *vs.* 2.8 ±0.5 BLE/min in noninfected animals; pairwise Holm’s adjusted *P* = 0.838). Similarly, infection reduced locomotion in YA animals by 12.5% (3.4 ±0.3 BLE/min in infected animals *vs.* 3.9 ±0.2 in noninfected animals; adjusted *P* = 0.788).

A multi-level meta-analysis combining the six infection-effect estimates from the four experiments showed that OrV infection had a significant negative effect on *C. elegans* motility (Fig. 1F: Hedges’ *g* = −0.667 ±0.160, *z* = −4.174, *P* < 0.001). Heterogeneity among experiments was negligible and not significant (τ^2^ = 4.23 ×10 ^−11^, *I*^2^ = 2.66×10^−8^; *Q* = 3.332, 5 d.f., *P* = 0.649), with all six effect size estimates being negative, indicating a consistent direction of effect across experiments (AGR.motility for YA: −0.61; PP.motility for L3: −0.65; SC.motility for L3: −1.45; SC.motility for YA: −0.76; Bİ.motility for L3: −0.28; and Bİ.motility for YA: −0.52).

In summary, across four independent motility assays and two developmental times, OrV infection consistently reduced *C. elegans* dispersal rates, supporting the conclusion that OrV impairs the movement of the host. The low estimated heterogeneity further suggests that this effect is robust across developmental stages and experimental conditions.

### OrV infection alters food choice in a cue-dependent manner

Fig. 2 summarizes the food-choice assays; Fig. 2A shows the experimental design. In the AGR.choice experiment, animals were given 1 h to make a choice between two drops of food. The probability of reaching either food patch was higher in YA than in L3 animals, but the age-related pattern differed by previous infection status (Fig. S1A). In L3, the estimated probability of having made a choice was 0.101 (95% CI: [0.053 - 0.183]) for noninfected animals and 0.350 [0.2576 - 0.456] for previously infected animals. Thus, at this time point, previous infection was associated with substantially greater odds of making a choice (OR = 4.815 [2.139 - 10.835], *P* < 0.001). In YA, the estimated probabilities of making a choice were 0.841 [0.705 - 0.922] for noninfected animals and 0.711 [0.546 - 0.834] for infected animals, and the difference between infection groups was not statistically significant (OR = 0.464 [0.165 - 1.302], *P* = 0.145). Accordingly, infection status and age interacted significantly in determining the probability of reaching either food patch (χ^2^ = 12.156, 1 d.f., *P* < 0.001), indicating that previous infection was associated with an increased probability of having made a choice at L3, rather than with a consistently greater probability of choosing across both developmental stages.

**FIG 2.**
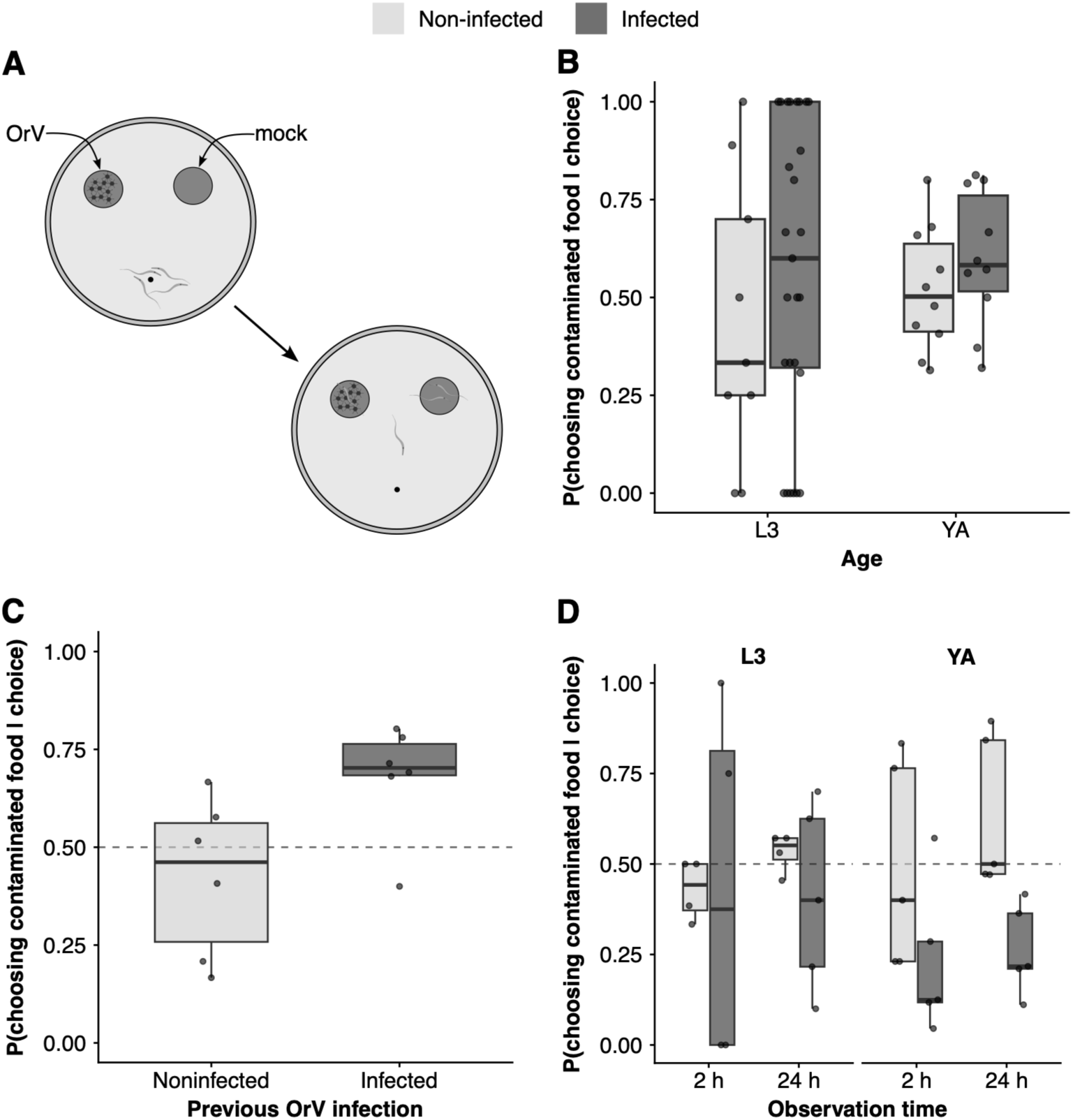
Effect of OrV infection on *C. elegans* food-choice behavior. **(A)** Schematic of the experimental design used to evaluate food choice. The same design was used in all experiments. **(B)** Experiment AGR.choice; probability that an animal that has moved toward food chooses the OrV-contaminated one depending on its infection status. Each dot represents an independent plate. **(C)** Experiment PP.choice; same as in panel (A), but observing only 24 h post-plating. **(D)** Experiment Bİ.choice; as in panel (B), but exploring the effect of developmental stage and whether or not food was OrV-contaminated on feeding behavior.

Among animals recovered from either food patch, however, there was no evidence of preference for contaminated or clean food in any experimental group (Fig. 2B). The estimated probabilities of choosing contaminated food were 0.453 [0.277 - 0.642] for noninfected L3 animals, 0.515 [0.386 - 0.641] for noninfected YA animals, 0.538 [0.415 - 0.656] for infected L3 animals, and 0.588 [0.452 - 0.712] for infected YA animals. All confidence intervals included the null expectation of 0.5, and none of the estimated probabilities differed significantly from an equal choice between the two food sources (all Holm’s adjusted *P* ≥ 0.807). Previous infection did not significantly affect the relative probability of choosing contaminated food at either L3 (OR = 1.404 [0.561 - 3.512], *P* = 0.469) or YA (OR = 1.349 [0.632 - 2.879], *P* = 0.439). Likewise, the probability of choosing food containing OrV did not change significantly between L3 and YA in either noninfected (OR = 1.280 [0.504 - 3.250], *P* = 0.604) or infected animals (OR = 1.230 [0.586 - 2.580], *P* = 0.585). There was also no evidence that the effect of infection on food preference differed between developmental stages, as indicated by the not significant infection × age interaction (χ^2^ = 0.004, 1 d.f., *P* = 0.948). Overall, previous OrV infection affected the probability of making a choice within the observation period but did not detectably alter which food patch was selected.

In the PP.choice experiment, previous OrV infection did not significantly affect the probability that L3 animals made a choice by 24 h post-exposure (Fig. S1B), with Beta-Binomial estimated choice probabilities of 0.632 [0.457 - 0.779] for noninfected and 0.680 [0.512 - 0.811] for infected animals, respectively. The odds of making a choice did not differ significantly between infected and noninfected animals (OR = 1.234 [0.460 - 3.312], *P* = 0.677). Among animals recovered from either food patch, previous infection significantly affected food preference (Fig. 2C; χ^2^ = 10.099, 1 d.f., *P* = 0.002). Noninfected animals showed no evidence of a preference for either food source: estimated probability of 0.405 [0.287 - 0.535], which did not differ significantly from the null expectation of 0.5 (*P* = 0.152). In contrast, infected animals chose contaminated food with an estimated probability of 0.682 [0.572 - 0.775], which was significantly greater than 0.5 (*P* = 0.003). Accordingly, previous infection increased the odds of choosing OrV-contaminated rather than clean food by approximately threefold relative to noninfected animals (OR = 3.146 [1.551 - 6.378], *P* = 0.002). These results indicate that previous OrV infection affected the direction of food choice, and was associated with greater selection of contaminated food, but did not detectably alter the overall probability that animals made a choice by 24 h post-exposure.

To distinguish responses to OrV from responses to other filtrate components, in the Bİ.choice experiment, we used a filtrate of healthy-animal as the mock preparation. Food choice behavior differed according to infection status, developmental stage, and observation time (early choice, 2 h, and late choice, 24 h). Among animals that reached a food patch (Fig. S1C), OrV infection significantly affected food preference overall (χ^2^ = 4.429, 1 d.f., *P* = 0.035), whereas neither developmental stage (χ^2^ = 0.277, 1 d.f., *P* = 0.598) nor observation time (χ^2^ = 1.558, 1 d.f., *P* = 0.212) had a significant overall effect. There was no statistical evidence that age modulated the effect of infection, as the infection × developmental stage interaction was not significant (χ^2^ = 2.196, 1 d.f., *P* = 0.138); this conclusion was also supported by direct comparisons of the infection effect between ages at 2 h (*P* = 0.198) and 24 h (*P* = 0.189). Furthermore, the effect of infection did not vary significantly with observation time (infection × time: χ^2^ = 1.508, 1 d.f., *P* = 0.219), and the three-way infection × developmental stage × time interaction was not significant (χ^2^ = 0.068, 1 d.f., *P* = 0.794). Nevertheless, the simple contrasts indicated that infected young adults had lower odds of choosing contaminated food than noninfected young adults at both 2 h (OR = 0.254 [0.073 - 0.878], *P* = 0.030) and 24 h post-exposure (OR = 0.155 [0.046 - 0.521], *P* = 0.003). No infection effect was detected in L3 larvae at either 2 h (OR = 1.101 [0.171 - 7.091], *P* = 0.919) or 24 h (OR = 0.517 [0.138 - 1.943], *P* = 0.329). When each experimental group was tested against the null expectation of equal choice between contaminated and clean food (Fig. 2D), only infected young adults at 2 h showed significant avoidance of contaminated food (OR = 0.238 [0.095 - 0.599], *P* = 0.018); the corresponding effect at 24 h did not remain significant after correction for eight tests (OR = 0.326 [0.136 - 0.782], *P* = 0.084). Overall, these results support an effect of OrV infection on food choice, with infected animals selecting the contaminated preparation less often, although the assay cannot distinguish avoidance of the preparation from attraction to healthy-animal cues. Although this effect was most evident in young adults, the interaction tests did not provide sufficient evidence to conclude that animal age statistically modulated the effect of infection.

Taken together, the first two experiments, infected animals appeared to prefer OrV-contaminated food. However, replacing food alone with food supplemented with healthy-animal filtrate reversed this pattern, with infected animals avoiding OrV-contaminated food. These findings suggest that OrV-clean food is more attractive than contaminated one, but that, when healthy-animal and infected-animal filtrates are compared directly, the healthy-animal filtrate is preferred. Therefore, OrV-associated food choice appears to reflect the combined and context-dependent effects of filtrate-derived and infection-specific cues.

### *eol-1* modulates infection-associated changes in locomotion and food choice

To determine whether *eol-1* contributes to infection-associated differences in locomotion, we compared the distances traveled over 30 minutes (SC.motility) by WT and *eol-1* animals at L3 and YA developmental stages under noninfected and infected conditions following the experimental design presented in Fig. 1A (Fig. 3A; the WT data are the same as those shown in Fig. 1D). Locomotion differed significantly among strains (*F*_1,69_ = 47.335, *P* < 0.001) and infection conditions (*F*_1,69_ = 16.872, *P* < 0.001), but not between developmental stages (*F*_1,69_ = 0.180, *P* = 0.673). Importantly, the effect of infection depended on genotype, as indicated by a significant strain × infection interaction (*F*_1,69_ = 6.783, *P* = 0.013). At the L3 stage, noninfected *eol-1* animals (3.0 ±0.9 BLE/min) moved 60.5% slower than noninfected WT animals (7.5 ±0.8 BLM/min; adjusted *P* < 0.001). The difference between infected *eol-1* animals (2.3 ±0.3 BLM/min) and infected WT animals (3.8 ±0.7 BLE/min) was smaller but remained significant (adjusted *P* = 0.043). Consistent with these observations, OrV infection markedly reduced locomotion in WT L3 animals (difference = −3.6 ±0.7 BLE/min, adjusted *P* < 0.001), whereas no significant effect of infection was detected in *eol-1* animals (difference = −0.7 ±0.7 BLE/min, adjusted *P* = 0.379). The difference between these infection effects was itself significant (difference = 3 ±1 BLE/min, adjusted *P* = 0.006). In YA animals, *eol-1* mutants also moved more slowly than WT animals under both noninfected (difference = −2.5 ±0.7 BLE/min, adjusted *P* = 0.002) and infected (difference = −1.6 ±0.7 BLE/min, adjusted *P* = 0.030) conditions. Infection did not significantly reduce motility in either WT YA animals (difference = −1.3 ±0.7 BLE/min, adjusted *P* = 0.082) or *eol-1* YA animals (difference = −0.5 ±0.7 BLE/min, adjusted *P* = 0.541). The three-way interaction among strain, infection status, and developmental stage was not significant (*F*_1,69_ = 2.028, *P* = 0.159). Together, these results indicate that *eol-1* strongly influences locomotion and may attenuate the reduction in movement associated with OrV infection. The evidence for this modulatory effect was strongest in L3 animals, in which the infection-induced decrease in motility observed in WT animals was largely absent in the *eol-1* mutant background.

**FIG 3.**
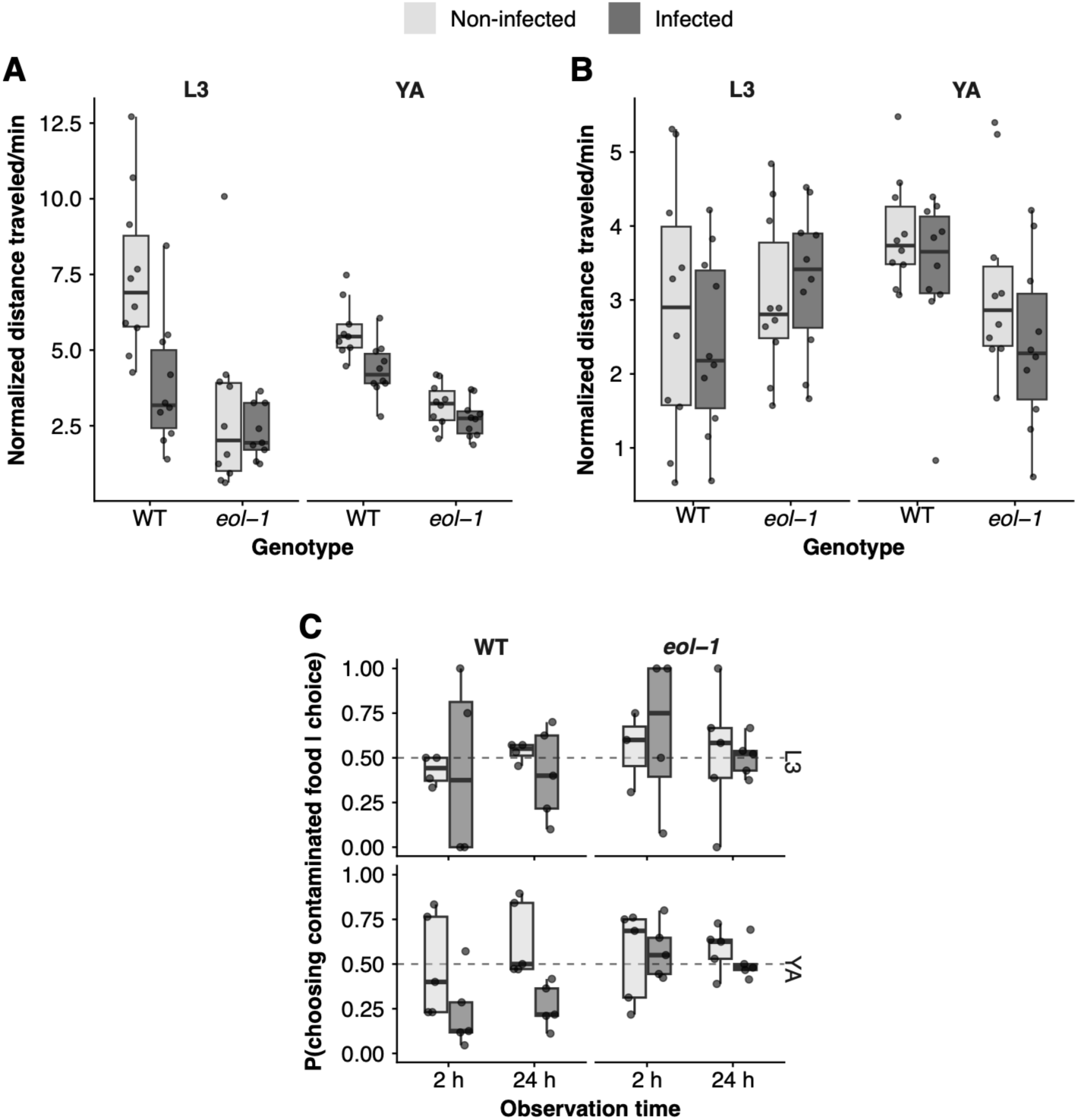
Effect of *eol-1* on motility and food choice. **(A)** Experiment SC.motility; effect of *eol-1* on the distance traveled in 30 min by noninfected and OrV infected L3 and YA animals. WT data are the same as those in Fig. 1D. **(B)** Experiment Bİ.motility; test of L3 and YA developmental stages on *eol-1* effect on motility after 65 min. WT data are the same as those in Fig. 1E. **(C)** Experiment Bİ.choice; test of the interaction between larval development and *eol-1* expression in the probability that noninfected and OrV infected animals chose OrV contaminated food. WT data are the same as those in Fig. 2D. In all panels, boxes indicate IQR, the horizontal line marks the median, and whiskers extend to 1.5×IQR beyond the quartiles.

We repeated the comparison of WT and *eol-1* animals over a 65 min observation period (Bİ.motility; Fig. 3B; the WT data are the same as those shown in Fig. 1E). Neither developmental stage (*F*_1,71_ = 1.798, *P* = 0.184), strain (*F*_1,71_ = 0.467, *P* = 0.497), nor infection status (*F*_1,71_ = 2.154, *P* = 0.147) had a significant overall effect on locomotor rate. However, the effect of developmental stage differed between strains, as indicated by a significant strain × stage interaction (*F*_1,71_ = 7.587, *P* = 0.008). At the L3 stage, *eol-1* did not significantly affect locomotion under either noninfected conditions (difference = −0.2 ±0.5 BLE/min, *P* = 0.719) or infected conditions (difference = 0.9 ±0.5 BLE/min, *P* = 0.092) relative to WT. In contrast, among YAs, *eol-1* animals tended to move more slowly than WT animals under both noninfected conditions (difference = −0.7 ±0.5, BLE/min, *P* = 0.159) and infected conditions (difference = −1.0 ±0.5 BLE/min, *P* = 0.048), although only the latter difference reached statistical significance. Infection had no detectable effect on locomotion in WT L3 animals (difference = −0.4 ±0.5 BLE/min, *P* = 0.386), *eol-1* L3 animals (difference = 0.2 ±0.5 BLE/min, *P* = 0.636), WT YAs (difference = −0.5 ±0.5 BLE/min, *P* = 0.333), or *eol-1* YA (difference = −0.8 ±0.5 BLE/min, *P* = 0.123). Consistent with these comparisons, the overall strain × infection interaction was not significant (*F*_1,71_ = 0.144, *P* = 0.705), and the effect of infection did not differ significantly between strains in either L3 animals (difference of differences = 0.7 ±0.7 BLE/min, *P* = 0.344) or YAs (difference of differences = −0.3 ±0.7 BLE/min, *P* = 0.679). Neither the infection × stage interaction (*F*_1,71_ = 1.145, *P* = 0.288) nor the three-way strain × infection × stage interaction *F*_1,71_ = 0.937, *P* = 0.336) was significant. Taken together, these results indicate that developmental stage influenced locomotor performance differently in WT and *eol-1* animals. However, there was no statistical evidence that *eol-1* altered the effect of OrV infection on motility over a 65 min observation period.

To further investigate the role of *eol-1* in food choice behavior, we extended the Bİ.choice experiment to include both infected and noninfected animals at the L3 and YA stages. The probability of making a choice depended strongly on developmental stage (Fig. S1D; χ^2^ = 21.391, 1 d.f., *P* < 0.001) and observation time (χ^2^ = 33.631, 1 d.f., *P* < 0.001). These effects were context dependent, as shown by significant strain × developmental stage (χ^2^ = 4.670, 1 d.f., *P* = 0.031), strain × time (χ^2^ = 26.590, 1 d.f., *P* < 0.001), strain × infection × developmental stage (χ^2^ = 4.655, 1 d.f., *P* = 0.031), strain × infection × time (χ^2^ = 48.787, 1 d.f., *P* = 0.003), infection × developmental stage × time (χ^2^ = 5.797, 1 d.f., *P* = 0.016), and strain × infection × developmental stage × time interactions (χ^2^ = 6.923, 1 d.f., *P* = 0.009). Thus, the influence of genotype and infection on whether animals made a choice varied jointly with developmental stage and observation time. Among animals that made a choice (Fig. 3C), there were no overall effects of strain (χ^2^ = 2.991, 1 d.f., *P* = 0.084), infection (χ^2^ = 2.168, 1 d.f., *P* = 0.141), developmental stage (χ^2^ = 0.030, 1 d.f., *P* = 0.862), or time (χ^2^ = 0.231, 1 d.f., *P* = 0.631) on the direction of food choice. Nevertheless, the strain × infection interaction was significant (χ^2^ = 3.929, 1 d.f., *P* = 0.047), indicating that the effect of OrV infection differed between WT and *eol-1* animals. This pattern was most evident in YA. Infection reduced the odds that WT YA chose contaminated food at both 2 h post-exposure (OR = 0.258 [0.086 - 0.772], *P* = 0.015) and 24 h (OR = 0.160 [0.055 - 0.462], *P* < 0.001), whereas no infection effect was detected in *eol-1* YA at either 2 hpi (OR = 0.984 [0.365 - 2.651], *P* = 0.974) or 24 hpi (OR = 0.839 [0.311 - 2.268], *P* = 0.730). Accordingly, infected *eol-1* YA had greater odds of choosing contaminated food than infected WT YA at 2 h (OR = 5.363 [1.809 - 15.900], *P* = 0.002) and 24 h (OR = 3.185 [1.121 - 9.050], *P* = 0.030). The strain-dependent effect of infection was formally supported in young adults at 24 hpi, where the ratio of ORs was 5.250 ([1.227 - 22.500], *P* = 0.025), but not at 2 h after exposure (ratio of Ors = 3.810 [0.869 - 16.700], *P* = 0.076). No comparable effects were detected in L3, and the higher-order interactions involving developmental stage and time were not significant for food preference. When individual groups were compared with an equal-choice probability of 0.5, only infected WT YA at 2 h post-exposure showed significant avoidance of OrV-contaminated food, with an estimated probability of 0.194 ([0.096 - 0.354], *P* = 0.011); the analogous tendency at 24 h post-exposure did not remain significant after correction for 16 tests (estimated probability = 0.247 [0.131 - 0.415], adjusted *P* = 0.069).

Overall, the Bİ.choice experiment (Fig. 3C) indicates that loss of *eol-1* disrupts the infection-associated avoidance of contaminated food observed in WT young adults.

Together, these results identify *eol-1* as a potential mechanistic contributor to the motility and behavioral changes induced by OrV infection, consistent with its infection-dependent overexpression and its known role in sensory regulation. Loss of *eol-1* attenuates the infection-associated reduction in motility and alters food choice behavior, indicating that OrV may impair dispersal, at least partly, through *eol-1*-dependent modulation of olfactory and locomotor responses.

## DISCUSSION

Across the four complementary assay designs, estimates of the infection effect estimates in the ERT54 background consistently indicated lower dispersal in OrV-exposed animals. The magnitude and statistical support of this association varied among experimental designs and was weakest in the 65 min window experiment, when neither stage-specific contrast was significant. Nevertheless, a multilevel meta-analysis supported an overall reduction in locomotion associated with infection.

Reduced dispersal is consistent with the sleep-like quiescent state reported during OrV infection (13); however, the present assays cannot distinguish slower movement from longer or more frequent quiescent bouts. Normalizing distance traveled is important because absolute displacement partly reflects the nearly twofold difference in body length between the L3 and YA developmental stages (37). OrV-induced quiescence of RNAi deficient animals has reduced-responsiveness and rapid-reversibility properties, is regulated by the ALA and RIS neurons, and promotes survival without measurably reducing viral load, suggesting that it contributes to infection tolerance or resilience rather than antiviral resistance (13). Expressing distance as the number of body-length equivalents traveled per minute therefore evaluates locomotor performance relative to body size rather than absolute displacement. Under this size-adjusted metric, developmental stage influenced the strain-dependent pattern of locomotor performance but did not demonstrably modify the effect of infection.

Reduced dispersal should not be assumed to be solely a nonspecific consequence of tissue damage. Infection-associated behavioral changes can be regulated components of host physiology. In bacterial systems, ingestion of pathogenic bacteria induces serotonin-dependent aversive learning (10,12,14), while microbial colonization and intestinal distension activate coordinated immune and neuroendocrine responses that promote pathogen aversion (38).

The food-choice experiments showed a context-dependent pattern, by which choice depended greatly on the alternative food source provided. Previous infection did not generate a context-independent preference for, or avoidance of, the contaminated preparation. Moreover, the probability of reaching either food patch must be distinguished from patch identity: failure to reach a patch can reflect reduced dispersal, altered exploration, or delayed arrival and should not be interpreted as sensory indecision. Among animals recovered from a patch, the direction of food choice depended strongly on how the alternatives were constructed. Such context dependence may explain the different outcomes among assays. When the contaminated patch uniquely contained material derived from infected individuals, animals could respond to viral particles, bacterial differences, animal metabolites, or other components of the preparation. Such context dependence is consistent with bacterial food-choice studies in which different components of a pathogenic bacterial preparation can have distinct, and sometimes opposing, behavioral effects. Defined bacterial metabolites can directly trigger avoidance through chemosensory neurons, as reported for *S. marcescens* and the *P. aeruginosa* metabolites phenazine-1-carboxamide and pyochelin (39,40). Conversely, volatile compounds associated with pathogenic *S. marcescens* can be attractive before or in the absence of the physiological consequences that drive learned avoidance (10,41). Infection can also modify the valence assigned to nonmicrobial chemical signals. In *P. aeruginosa*-infected hermaphrodites, induction of the STR-44 chemoreceptor in AWA neurons changes the response to an ascaroside pheromone mixture and promotes mating, demonstrating that infection state can reconfigure the sensory interpretation of an otherwise familiar external cue (42). Consequently, attraction to a component of an infected-animal preparation and avoidance learned through infection need not produce the same choice direction. By contrast, the Bİ.choice design matched animal-derived cues between the alternatives by comparing filtrates from healthy and infected animals. Under these conditions, infected YA selected the OrV-containing patch less frequently. Therefore, we interpret Bİ.choice as the most specific test of an infection-associated food response, while recognizing that complete chemical equivalence between preparations cannot be assumed.

These results do not support a single pooled food choice effect across all designs. Instead, they indicate that OrV-associated selection is cue-dependent and can change direction when animal-derived cues are not matched. The contrasting food choice outcomes among assay designs may therefore be biologically informative rather than merely reflecting experimental inconsistency because each design presented a different combination of viral, bacterial, and animal-derived cues. Claims about attraction to or avoidance of OrV should therefore be restricted to designs that isolate viral associations from other components of the inoculum. The dependence of choice on both prior infection and cue composition suggests that the response may involve an interaction between internal infection state and external sensory information, rather than invariant attraction to or avoidance of a viral component. In bacterial systems, *C. elegans* integrates directly detected microbial metabolites with infection-dependent serotonergic and neuroendocrine signals, allowing prior experience and physiological state to change the valence of food-associated cues (10,38,40).

The *eol-1* experiments support a genetic contribution but did not establish a complete mechanism. Loss of *eol-1* strongly altered baseline dispersal in the Bİ.motility assay. The genotype × infection interaction and planned contrasts provided the clearest evidence for an EOL-1 contribution to the infection-associated reduction in dispersal of L3 animals. However, because the strain × infection × time interaction was not significant, the data do not formally demonstrate that this contribution was restricted to the earlier observation time. The absence of a detectable genotype × infection interaction in the 65 min experiment nevertheless indicates that EOL-1 was not universally required for the dispersal phenotype under all experimental conditions.

A similar genotype-dependent pattern emerged in the metabolite-controlled food-choice assay. Infection reduced the selection of OrV-containing food among WT young adults, whereas the corresponding infection contrasts were not significant in *eol-1* mutants. The overall strain × infection interaction supports a contribution of EOL-1 to the food-choice response, and the age- and time-specific contrasts indicate that this pattern was clearest in young adults. However, the not significant higher-order interactions do not demonstrate that the EOL-1-dependent infection effect was restricted to a particular developmental stage or observation time. Because *eol-1* acts in URX sensory neurons to constrain experience-dependent changes in olfactory preference for bacterial food (33), the present results raise the possibility that EOL-1 influences the integration of internal infection state with external food-associated cues. This interpretation remains provisional because the mutant also displayed a baseline locomotor phenotype, and the experiments did not include neuronal rescue or direct measurements of sensory-neuron activity. Another possibility is that EOL-1 is acting earlier on, during the generation of the antiviral RNAi response (36), and that this response is necessary for neuroimmune signaling and behavioral changes.

In conclusion, our experiments support an overall association between OrV infection and reduced *C. elegans* dispersal. A multilevel synthesis indicated a common negative direction across assay designs, but the magnitude and statistical support of this association varied substantially among experimental contexts. After accounting for body size, there was no evidence that the effect of infection differed between developmental stages. The effect of infection on food selection depended on the composition of the alternatives, emphasizing the need to distinguish responses to virus-associated cues from responses to other components of animal-derived preparations. EOL-1 contributed to the early dispersal response and modified infection-associated food choice; however, the genetic analyses did not establish a specific neuronal or molecular mechanism. Future experiments combining continuous movement tracking, individual viral-load measurements, chemical characterization of matched filtrates, and cell-specific rescue of *eol-1* will be needed to connect infection state, neural processing, and behavior.

## MATERIALS AND METHODS

### *C. elegans* strains and growth conditions

Animals were cultured and maintained at 20°C on nematode growth medium (NGM) agar plates seeded with *Escherichia coli* OP50 (43). The transgenic strain ERT54 (*jyIs8[pals-5p::GFP + myo-2p::mCherry]X*), which carries a wild-type Bristol N2 genetic background and expresses GFP in response to intracellular infection (44), was used as a surrogate wild-type.

Strain SFE10 (*eol-1(tm6609), jyls8[pals-5p::GFP + myo-2p::mCherry]X*) was used for testing the role of sensory processing in OrV-induced effects in locomotion. SFE10 and ERT54 are isogenic with the exception of the *eol-1* mutation in the former.

To synchronize populations, plates containing embryos were gently washed with M9 buffer (0.22 M KH_2_PO_4_, 0.42 M Na_2_HPO_4_, 0.85 M NaCl, 1 mM MgSO_4_) to remove larvae and adults while leaving the embryos behind. After 1 h, plates were washed again with M9 buffer to collect larvae that had hatched during that interval, and these were transferred to freshly seeded NGM plates.

### OrV isolate, stocks, and inoculation conditions

OrV stock was generated by inoculating two 6 cm plates of freshly starved individuals of the more susceptible strain JU2624 (*mjIs228[myo-2::mcherry::unc54; lys-3p::eGFP::tbb-2]*, JU1580 background) with the JUv1580_vlc (45) OrV strain and expanding them on fifty 9 cm NGM plates until starvation. Plates were washed with M9 buffer and pooled into 15 mL tubes. Tubes were vortexed for 30 s, incubated at room temperature for 10 min, and centrifuged at 2,250×g for 2 min. The animal pellet was discarded, and the supernatant was transferred to a new tube, centrifuged twice at 21,000×g for 5 min at 4 °C, and filtered through a 0.22 μm syringe filter. The resulting viral preparation was stored at −80 °C.

Viral RNA extraction and quantification using RT-qPCR have been described elsewhere (32,36,45–47). Briefly, RNA was extracted using the Viral RNA Isolation Kit (NZYtech, Lisbon, Portugal) following the manufacturer’s instructions. OrV titers were quantified by standard curve RT-qPCR using the qPCRBIO SyGreen 1-Step Go Hi-ROX kit (PCR Biosystems Ltd., London, UK) on an ABI StepOnePlus Real-Time PCR System (Applied Biosystems, Foster City CA, USA), with primers targeting a region of OrV RNA2 (5’-ACGAAGCAGTAGCCGTTAAG-3’ and 5’-GAGAACATCCTTCTCTGCGG-3’).

Larvae were always inoculated at hatching by applying OrV stock containing a total of 2.8×10^9^ copies of RNA2 onto *E. coli* OP50 lawns. This dose was used because it infects close to 100% of animals at 24 hours post-inoculation (32). Control plates were treated with an equivalent volume of M9 buffer.

### Dispersal assays and image processing

For all individual-level dispersal assays, distance traveled was divided by the median body length corresponding to the developmental stage tested and is therefore reported as the number of body lengths traveled (48). Therefore, distances are expressed as body-length equivalents (BLE).

In the first experiment (design AGR.motility), either 23 ±24 (±1.5×IQR) noninfected or 23 ±7 infected YA (48 hours post-hatching) were placed in the center of a 6 cm diameter seeded plate. Five concentric circles with 0.3 cm intervals were drawn. After 48 h, the number of animals in each of the six defined areas was recorded. Seven replicates per treatment were performed.

In the second set of experiments (design PP.motility), 3.5 cm diameter plates, with *E. coli* OP50 covering the entire plate surface, were seeded with a single animal in the third larval stage (L3, 24 hours post-hatching) in the center and allowed to move away. Eleven replicates for noninfected animals and ten replicates for infected animals were performed. After 105 min, pictures were taken using a mobile phone camera, analyzed using FIJI (49), and distance traveled was measured by drawing the path followed by each individual animal, twice per path.

In the third experiment (design SC.motility), the traveled distance was measured after 30 min as in experiment PP.motility, but developmental stages L3 and YA were used. Because this design compared L3 and YA, stage-specific median body lengths of 0.43 ±0.12 mm (±1.5×IQR) and 0.82 ±0.21 mm (data from (36)), respectively, were used to normalize distances traveled.

In the fourth experiment (design Bİ.motility), the traveled distance was measured as in experiments PP.motility and SC.motility, but observations were made 65 min after single animals were placed in the center of the plates. Both L3 and YA animals were used with 10 replicates per developmental stage and inoculation status.

To test whether *eol-1* contributes to the observed difference in motility, we used *eol-1(tm6609)* animals and measured motility as in experiments SC.motility and Bİ.motility.

### Food-choice assays

Here, food choice to refers to the tendency of animals to move toward OrV-contaminated food *vs*. uncontaminated food. Synchronized populations were washed from plates using M9 buffer and left to sediment by gravity for 15 minutes. Animals were then pipetted on a designated point without food in 6 cm diameter plates. Two food drops were placed diagonally at equal distances from the starting point. One spot contained mock food (M9 or healthy JU1580 animal filtrate), and the other contained food mixed with 20 μL of OrV filtrate (3.32×10^8^ copies of RNA2). Four experimental blocks, differing in details, were performed.

In the first block (design AGR.choice), the experiment was performed with noninfected and infected L3 and YA animals, with 10 replicates per treatment and a median of 39 (18 IQR) animals per plate. After 1 h, the number of animals on each drop and elsewhere on the plate was counted.

The second block (design PP.choice) followed the same design, except the plates were observed 24 h after plating the animals. We performed six replicates each with 12 ±10 (1.5×IQR) noninfected L3 animals and 11 ±9 infected L3 animals per plate.

The third block (design Bİ.choice) was performed using both infected and noninfected L3 and YA animals. The main difference between this design and the previous ones was that the mock food spot was diluted into a filtrate from healthy animals, prepared as described for OrV stock preparation above, in the same proportion filtrate:bacteria as in the case of OrV-contaminated food. In this case, five replicates per time point and developmental stage and infection condition, with a median of 17 ±14 (1.5×IQR) animals per plate.

To test whether *eol-1* contributes to the observed difference in food choice, *eol-1(tm6609)* animals were used (Bİ.choice as described above).

### Statistical analyses

To formally test the effect of infection on dispersal in the AGR.motility experiment, the observed distribution of animals across ordinal distance classes was analyzed using a cumulative link mixed model (CLMM). Infection status was included as a fixed effect, whereas plate identity was included as a random effect to account for variation among replicate plates. CLMM was fitted using R package ‘ordinal’ version 2026.7-26.

Dispersal data generated with the experimental design PP.motility assays were fitted to a GLMM with a Gaussian error distribution and identity-link function. Infection status was included as a fixed effect, and plate identity was added as a random effect to account for replication. For the design SC.motility, infection status, observation time, and their interaction were included as fixed effects. For the design Bİ.motility, infection status, developmental stage, and their interaction were included as fixed effects. In the analysis including ERT54 and *eol-1*, strain, infection status, developmental stage, and all corresponding interactions were included as fixed effects. In both cases, the generalized linear model (GLM) used a Gaussian error distribution and identity-link function.

A multilevel meta-analysis of locomotion across the four independent experiments was performed using standardized mean differences (Hedges’ *g*) and a random-effects model fitted by restricted maximum likelihood (REML) with the R package ‘metafor’ version 5.0-1. Effect sizes were calculated as the mean dispersal in the infected group minus that in the control group; therefore, *g* < 0 values indicate reduced dispersal following infection.

For food-choice responses in the AGR.choice design, plates examined at 24 and 48 h post-exposure were treated as independent observations and fitted to a GLM with a Binomial error distribution and logit-link function. For all other food-choice designs, data were analyzed using GLMMs with a Binomial or a Beta-Binomial distribution and a logit-link function, selecting the error structure by AIC. Infection status, observation time, and their interaction were included as fixed effects, and plate identity was included as a random effect to account for replication. The analyses separated two conditional components. First, food-patch arrival was analyzed by comparing the number of animals recovered from either food patch with the number recovered elsewhere on the plate. Second, patch identity among animals recovered from a food patch was analyzed by comparing the number selecting the OrV-containing preparation with the number selecting the clean preparation; plates with no animals in either patch were excluded from this component. Preference was evaluated by comparing the estimated probability of selecting the OrV-containing preparation with the null expectation of 0.5, whereas the effect of infection was evaluated by contrasting infected and noninfected animals within each observation time.

GLMs and GLMMs were fitted using the R package ‘glmmTMB’ version 1.1.14. Estimated marginal probabilities, 95% confidence intervals, and pairwise contrasts were obtained using the R package ‘emmeans’ version 2.0.3 with Holm adjustment for multiple comparisons.

All statistical analyses were performed using R version 4.6.0 under RStudio version 2026.09.0+174 (Posit PBC, Boston MA, USA). Plots were generated using ‘ggplot2’ version 4.0.3.

## SUPPLEMENTAL MATERIAL

**FIG S1.**
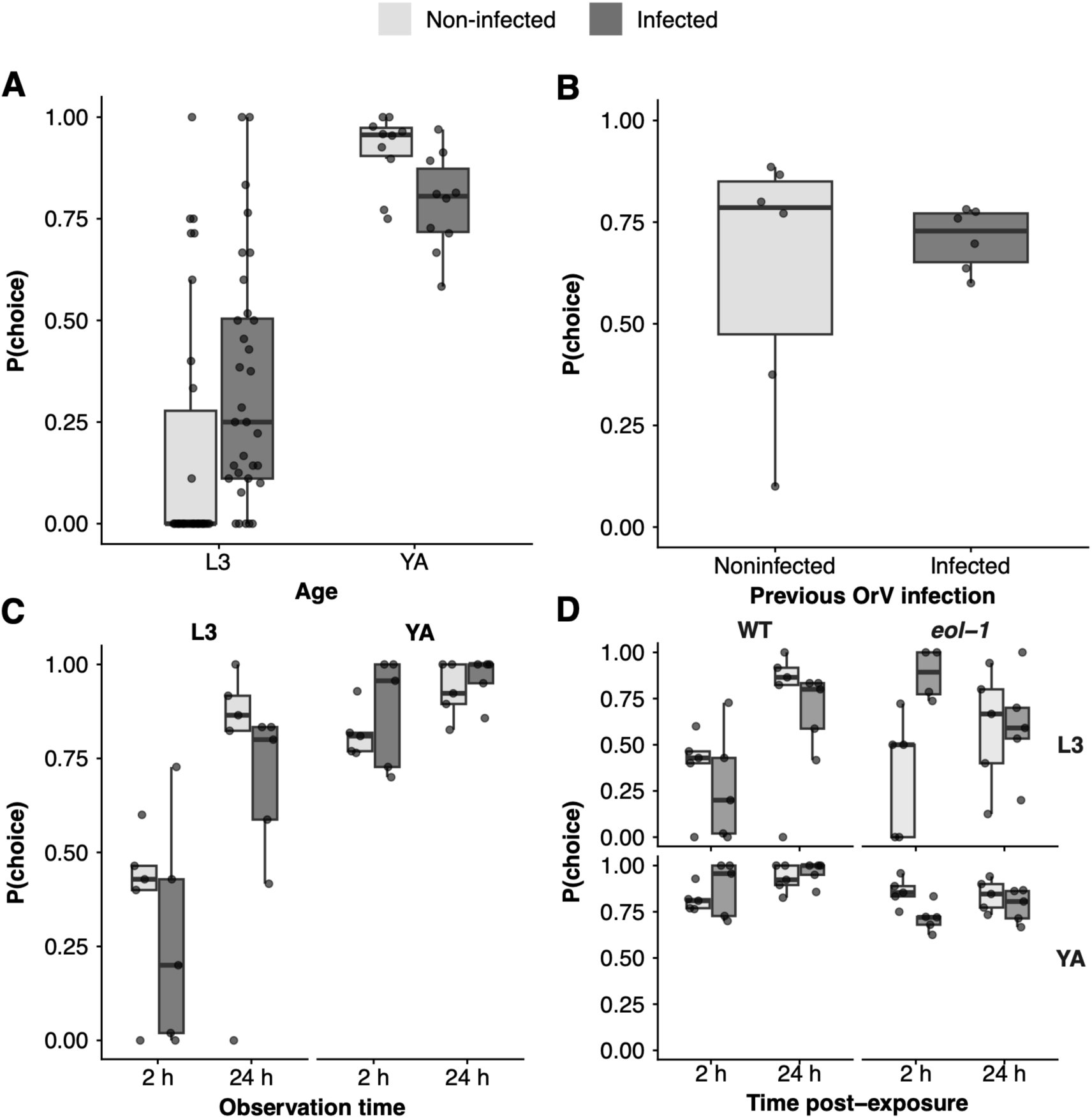
Effect of OrV infection in *C. elegans* initiation of movement toward food. **(A)** Experiment AGR.choice; probability that a animal moves toward food depending on its infection status. Each dot represents an independent plate. Plates were observed 1 h after exposure. **(B)** Experiment PP.choice; same as in panel (A) but observing only 24 h post-exposure. **(C)** Experiment Bİ.choice; as in panel (A) but exploring the effect of developmental stage in moving behavior. **(D)** Test of the interaction between larval development and *eol-1* expression in the probability that noninfected and OrV infected animals initiate movement toward food (experiment Bİ.choice; ERT54 data are the same than in Fig. S1C). In all panels, boxes indicate IQR, the horizontal line marks the median, and whiskers extend to 1.5×IQR beyond the quartiles.

## ETHICS APPROVAL

This study did not require ethical approval from a human subject or animal welfare committee.

## DATA AVAILABILITY

All data and R code accompanying the paper can be found on Zenodo (https://doi.org/10.5281/zenodo.23100300).

## ACKNOWLEDGMENTS

We thank Francisca de la Iglesia for her excellent technical support and the members of the EvolSysVir lab for their valuable comments and fruitful discussions. We also thank Animalbase and the Caenorhabditis Genetics Center.

This work was supported by grants PID2025-169610NB-I00, funded by MCIU/AEI/10.13039/501100011033 and by “ERDF a way of making Europe”, and CIPROM/2022/59, funded by Generalitat Valenciana to S.F.E. E.G.L. was supported by grant FPU21/00410 funded by MCIU/AEI/10.13039/501100011033 and by “ESF invest in your future”. D.H. was supported by grant PREP2022-000699 funded by MCIU/AEI/10.13039/501100011033 and by “ESF investing in your future”. V.G.C. was supported by grant MSCA 2024-PF-01-101207897 funded by Horizon Europe.

## AUTHOR’S CONTRIBUTION

E.G.L., D.H., V.G.C., and S.F.E. designed research; E.G.L., D.H., I.M., A.V.G., S.C., A.G.R., B.İ., and P.P. performed research; E.G.L., D.H., V.G.C., and S.F.E. supervised research; S.F.E. analyzed data; S.F.E. wrote the paper.

All authors gave final approval for publication and agreed to be held accountable for the work performed therein.

